# Visual self-motion cues are impaired in Parkinson’s disease yet over-weighted during visual-vestibular integration

**DOI:** 10.1101/2019.12.22.884940

**Authors:** Sol Yakubovich, Simon Israeli-Korn, Orly Halperin, Gilad Yahalom, Sharon Hassin-Baer, Adam Zaidel

**Author notes:** These authors contributed equally to this work (Sol Yakubovich, Simon Israeli-Korn). Corresponding author: Adam Zaidel Gonda Multidisciplinary Brain Research Center Bar Ilan University Ramat Gan, 52900 Israel.

## Abstract

**Background:** Parkinson’s disease (PD) is prototypically a movement disorder. Although perceptual and motor functions are interdependent, much less is known about perceptual dysfunction in PD. Perceptual deficits can impact activities of daily living, and contribute to motor symptoms, but might go unnoticed if not tested directly. Posture, gait and balance, affected in PD, rely on veridical perception of one’s own motion in space. Yet it is unknown whether self-motion perception is impaired in PD.

**Objectives:** To test self-motion perception in PD, separately for visual and vestibular cues (unisensory), and multisensory integration thereof.

**Methods:** Participants (19 early stage PD, 23 age-matched and 20 young adult controls) experienced vestibular (motion platform), visual (optic flow), and combined visual-vestibular self-motion stimuli, and discriminated whether the stimulus headings were rightward or leftward of straight ahead. PD participants and age-matched controls were tested on two visits (PD on and off medication).

**Results:** PD participants had significantly impaired visual self-motion perception, both on and off medication. This deficit correlated significantly with clinical disease severity. By contrast, their vestibular performance was unimpaired. Remarkably, despite impaired visual self-motion perception, PD participants significantly over-weighted visual cues during multisensory (visual-vestibular) integration.

**Conclusions:** Self-motion perception is affected already in early stage PD, specifically by impaired visual (vs. vestibular) function, and by suboptimal visual-vestibular integration. This may contribute to impaired balance and gait control. Future investigation into this connection might open up new avenues for alternative therapies to better treat these symptoms. Furthermore, these results may also impact early PD diagnosis and subtyping.

## INTRODUCTION

Parkinson’s disease (PD) is primarily characterized by a decline in motor function, marked by the cardinal features of bradykinesia, akinesia, rigidity, tremor and postural instability (Parkinson, 1817; Jankovic, 2008; Postuma *et al.*, 2015). Although James Parkinson already noted many of the non-motor aspects of PD in his original seminal description (Parkinson, 1817), these are only recently receiving more substantial attention (Chaudhuri *et al.*, 2006; Patel *et al.*, 2014). In contrast to motor symptoms, non-motor symptoms (such as perceptual deficits) are by nature less observable, under-reported and may go unnoticed if not actively and directly tested (Shulman *et al.*, 2002; Chaudhuri *et al.*, 2010; Bonnet *et al.*, 2012).

Veridical perception of one’s orientation and self-motion in space is fundamental for motor control, especially gait and balance (Dichgans and Brandt, 1978; Paillard, 1991; Horak and Macpherson, 1996; Lackner and DiZio, 2005; Cullen, 2012). Self-motion perception relies primarily on vestibular and visual (optic flow) cues (Dichgans and Brandt, 1978; Warren and Hannon, 1988; Fushiki *et al.*, 2005; Gu *et al.*, 2007; Fetsch *et al.*, 2009, 2010; Butler *et al.*, 2010; Zaidel *et al.*, 2015), as well as other somatosensory cues, such as proprioception (Probst *et al.*, 1985; Hlavačka *et al.*, 1992; Mergner *et al.*, 1993; Hlavacka *et al.*, 1996; Mergner and Rosemeier, 1998; Schweigart *et al.*, 2002; Durgin *et al.*, 2005). Furthermore, these unisensory cues need to be integrated to form a unified and reliable percept of self-motion (Angelaki *et al.*, 2009; Butler *et al.*, 2010). It is currently not known: i) whether or not vestibular/visual perception of self-motion is impaired in PD nor ii) whether multisensory integration is affected in PD. In this study, we directly tested unisensory vestibular and unisensory visual performance, as well as multisensory integration of visual and vestibular cues for self-motion perception in PD.

Vestibular deficits in PD are a matter of debate. Reichert et al. (1982) demonstrated altered nystagmus responses in PD to caloric stimulation. But, this could reflect sensory-motor integration deficits, rather than vestibular deficits *per se*. Lithgow and Shoushtarian (2015) also report altered vestibular responses in PD. But, the technique used in that study (electrovestibulography, which measures vestibular responses from the ear canal) is esoteric and somewhat controversial (Brown *et al.*, 2017). By contrast, Bertolini et al. (2015) did not find vestibular sensory impairment, but rather deficits that likely reflect faulty integration of vestibular signals in the brain. Thus, there is little evidence to support vestibular dysfunction in PD. However, this requires further validation (see Smith, 2018 for a review).

PD patients demonstrate altered navigational veering in response to visual self-motion (optic flow) stimuli (Davidsdottir *et al.*, 2008; Lin *et al.*, 2014), as well as reduced activation in visuomotor brain areas (van der Hoorn *et al.*, 2014). While these studies imply visual deficits of self-motion perception, they do not isolate whether or not there is a specific perceptual (vs. sensorimotor) deficit, because veering is a graded motor response. Hence, the need to perform an experiment that specifically tests visual self-motion perception.

A different type of visual motion stimulus, that is not designed to elicit a percept of self-motion, uses random dot kinematograms (RDKs). RDKs comprise dots moving in 2D, presented through an aperture on a flat screen in front of the observer, in order to test coherent perception of visual motion in the environment. PD Performance in RDK experiments did not differ from controls (Putcha *et al.*, 2014; Jaywant *et al.*, 2016). However, Putcha *et al*. (2014) did find an association between increased discrimination thresholds and disease severity. There are also many other visual impairments described in PD, including delays in visual evoked responses, abnormalities in contrast, spatiotemporal and color sensitivity and altered perception of visual orientation (Bodis-Wollner and Yahr, 1978; Bodis-Wollner *et al.*, 1987; Montse *et al.*, 2001; Gullett *et al.*, 2013; Weil *et al.*, 2016, 2017, 2018). Thus, visual motion perception in PD requires further investigation.

Intriguingly, despite these abovementioned visual perception deficits, PD patients seem to be functionally more dependent on vision (Cooke *et al.*, 1978; Bronstein *et al.*, 1990; Azulay *et al.*, 1999, 2002; Almeida and Lebold, 2010; Cowie *et al.*, 2010). Thus, in contrast to healthy humans (Jacobs, 1999; Landy and Kojima, 2001; Ernst and Banks, 2002; Alais and Burr, 2004; Fetsch *et al.*, 2009; Raposo *et al.*, 2012) and animals (Gu *et al.*, 2008; Raposo *et al.*, 2012) and even other clinical populations (tested in autism, Zaidel *et al.*, 2015), that largely follow Bayesian predictions of multisensory integration, we hypothesized that PD patients may have a specific (and perhaps unique) multisensory integration impairment, with visual overweighting. Here we tested this hypothesis directly, within the Bayesian framework of multisensory integration.

Using a multisensory motion simulator, we tested visual, vestibular, and combined (visual-vestibular) perception of self-motion in PD patients and controls. To avoid confounding effects of motor dysfunction, participants performed a binary two-alternative forced choice (2AFC) psychophysics task. Testing the same task (heading discrimination) with the same categorical responses for visual or vestibular cues, and in the same participants, allowed us to isolate specific visual and/or vestibular perceptual (dys)function. Strikingly, we found impaired visual self-motion perception in PD, whereas vestibular performance was unimpaired. Finally, by introducing a small discrepancy between the visual and vestibular cues (in the combined condition) we tested multisensory integration, and found that PD patients over-weighted the visual cues (under-weighting vestibular) despite the visual impairment, exposing sub-optimal multisensory integration.

## MATERIALS AND METHODS

### Participants and procedures

We tested 20 patients with (early stage) idiopathic PD, recruited through the Movement Disorders Institute at Sheba Medical Center, 24 age-matched controls (recruited from the general public, spouses of the PD participants, and staff at Bar Ilan University), and 21 young adult controls (recruited from the student body at Bar Ilan University). One participant from each group was excluded due to inadequate task performance as evidenced by close to arbitrary heading choices. This resulted in 19 PD, 23 age-matched and 20 young-adult participants, for further analysis. This study was approved by the Internal Review Boards of Bar Ilan University and Sheba Medical Center. All participants signed informed consent prior to partaking in the study and received compensation for participation. Exclusion criteria for recruitment to the study included: neurological or psychiatric conditions (apart from PD), inability to walk independently or to climb stairs safely unassisted, poor corrected vision, deafness, dementia or vestibular dysfunction. Cognitive function was assessed in all participants using the Montreal Cognitive Assessment (MoCA) test (Nasreddine *et al.*, 2005).

Individual participant details of the PD, age-matched and young adult groups are presented in Tables 1-3, respectively. The PD and age-matched groups had comparable age (mean ± SD = 62.9 ± 8.9 and 62.7 ± 6.9 years, respectively), gender (68% and 65% male, respectively) and cognitive function (mean MoCA score ± SD = 25.8 ± 3.1 and 24.5 ± 2.3, respectively). Disease severity in PD participants was measured according to the motor part of the Unified Parkinson’s Disease Rating Scale (UPDRS) and is reported in Table 1. Young controls were aged 23.9 ± 3.0 years, were 50% male and had MoCA scores 27.5 ± 1.5.

**Table 1:**
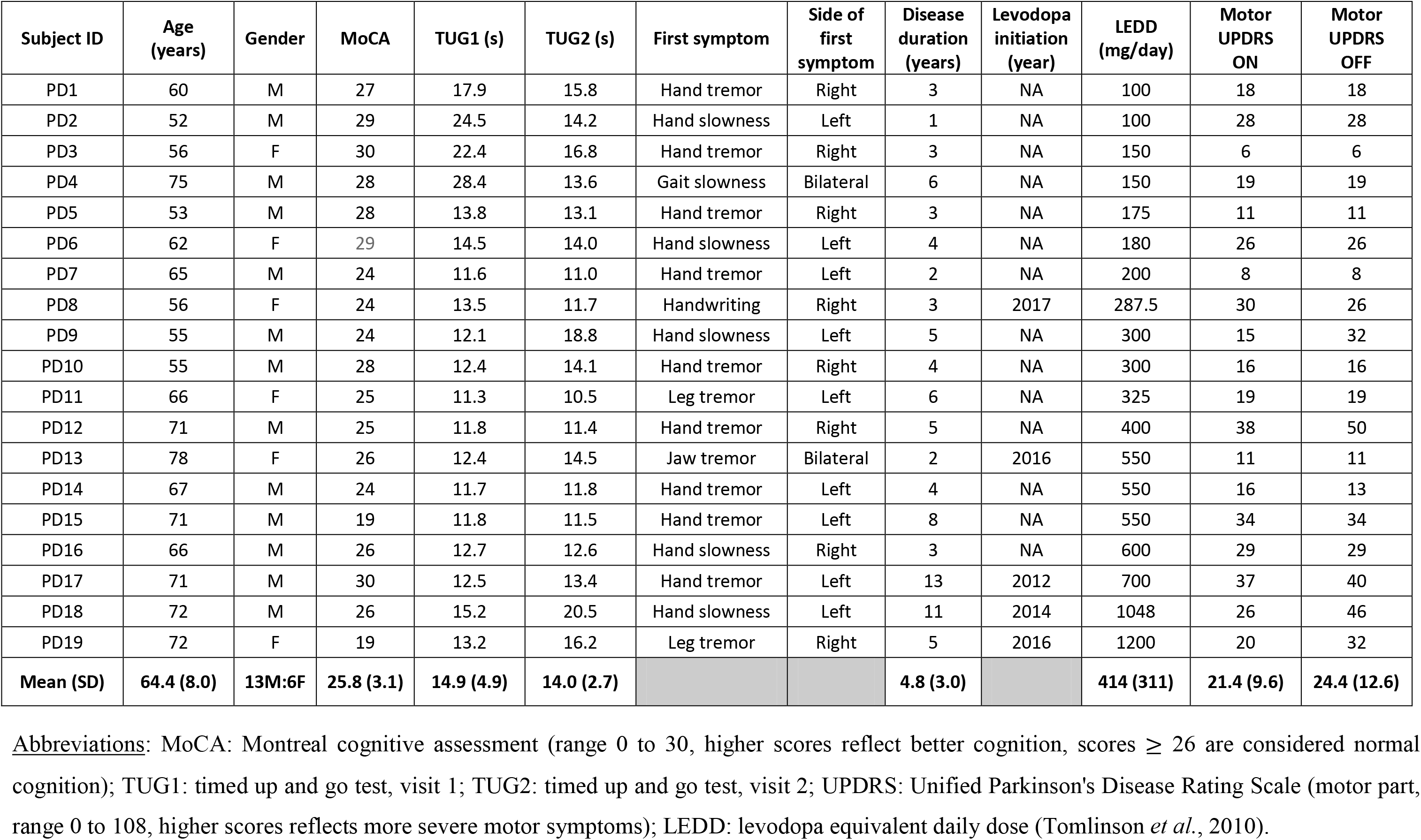
Parkinson’s disease participants’ details

| Subject ID | Age (years) | Gender | MoCA | TUG1 (s) | TUG2 (s) | First symptom | Side of first symptom | Disease duration (years) | Levodopa initiation (year) | LEDD (mg/day) | Motor UPDRS ON | Motor UPDRS OFF |
| --- | --- | --- | --- | --- | --- | --- | --- | --- | --- | --- | --- | --- |
| PD1 | 60 | M | 27 | 17.9 | 15.8 | Hand tremor | Right | 3 | NA | 100 | 18 | 18 |
| PD2 | 52 | M | 29 | 24.5 | 14.2 | Hand slowness | Left | 1 | NA | 100 | 28 | 28 |
| PD3 | 56 | F | 30 | 22.4 | 16.8 | Hand tremor | Right | 3 | NA | 150 | 6 | 6 |
| PD4 | 75 | M | 28 | 28.4 | 13.6 | Gait slowness | Bilateral | 6 | NA | 150 | 19 | 19 |
| PD5 | 53 | M | 28 | 13.8 | 13.1 | Hand tremor | Right | 3 | NA | 175 | 11 | 11 |
| PD6 | 62 | F | 29 | 14.5 | 14.0 | Hand slowness | Left | 4 | NA | 180 | 26 | 26 |
| PD7 | 65 | M | 24 | 11.6 | 11.0 | Hand tremor | Left | 2 | NA | 200 | 8 | 8 |
| PD8 | 56 | F | 24 | 13.5 | 11.7 | Handwriting | Right | 3 | 2017 | 287.5 | 30 | 26 |
| PD9 | 55 | M | 24 | 12.1 | 18.8 | Hand slowness | Left | 5 | NA | 300 | 15 | 32 |
| PD10 | 55 | M | 28 | 12.4 | 14.1 | Hand tremor | Right | 4 | NA | 300 | 16 | 16 |
| PD11 | 66 | F | 25 | 11.3 | 10.5 | Leg tremor | Left | 6 | NA | 325 | 19 | 19 |
| PD12 | 71 | M | 25 | 11.8 | 11.4 | Hand tremor | Right | 5 | NA | 400 | 38 | 50 |
| PD13 | 78 | F | 26 | 12.4 | 14.5 | Jaw tremor | Bilateral | 2 | 2016 | 550 | 11 | 11 |
| PD14 | 67 | M | 24 | 11.7 | 11.8 | Hand tremor | Left | 4 | NA | 550 | 16 | 13 |
| PD15 | 71 | M | 19 | 11.8 | 11.5 | Hand tremor | Left | 8 | NA | 550 | 34 | 34 |
| PD16 | 66 | M | 26 | 12.7 | 12.6 | Hand slowness | Right | 3 | NA | 600 | 29 | 29 |
| PD17 | 71 | M | 30 | 12.5 | 13.4 | Hand tremor | Left | 13 | 2012 | 700 | 37 | 40 |
| PD18 | 72 | M | 26 | 15.2 | 20.5 | Hand slowness | Left | 11 | 2014 | 1048 | 26 | 46 |
| PD19 | 72 | F | 19 | 13.2 | 16.2 | Leg tremor | Right | 5 | 2016 | 1200 | 20 | 32 |
| **Mean (SD)** | **64.4 (8.0)** | **13M:6F** | **25.8 (3.1)** | **14.9 (4.9)** | **14.0 (2.7)** |  |  | **4.8 (3.0)** |  | **414 (311)** | **21.4 (9.6)** | **24.4 (12.6)** |

Abbreviations: MoCA: Montreal cognitive assessment (range 0 to 30, higher scores reflect better cognition, scores $\geq 26$ are considered normal cognition); TUG1: timed up and go test, visit 1; TUG2: timed up and go test, visit 2; UPDRS: Unified Parkinson's Disease Rating Scale (motor part, range 0 to 108, higher scores reflects more severe motor symptoms); LEDD: levodopa equivalent daily dose (Tomlinson *et al.*, 2010).

**Table 2:** Age-matched control participants’ details

| Subject ID | Age (years) | Gender | MoCA | TUG1 (s) | TUG2 (s) |
| --- | --- | --- | --- | --- | --- |
| AMC1 | 62 | F | 25 | 11.7 | 10.9 |
| AMC2 | 49.5 | M | 26 | 10.7 | 11.7 |
| AMC3 | 64 | M | 22 | 11.9 | NA |
| AMC4 | 65 | F | 28 | 10.6 | 11.4 |
| AMC5 | 59 | F | 23 | 12.2 | 19.3 |
| AMC6 | 72 | M | 22 | 14.9 | 13.6 |
| AMC7 | 70 | M | 22 | 13.0 | 23.7 |
| AMC8 | 65.5 | M | 25 | 15.2 | 12.7 |
| AMC9 | 70 | F | 20 | 12.8 | 12.4 |
| AMC10 | 57 | M | 27 | 10.1 | 9.2 |
| AMC11 | 65 | M | 26 | 9.6 | 9.8 |
| AMC12 | 56 | F | 25 | 16.7 | 15.8 |
| AMC13 | 68 | F | 24 | 14.6 | 14.6 |
| AMC14 | 68.5 | M | 26 | 10.3 | 11.4 |
| AMC15 | 63 | M | 23 | 9.0 | 10.5 |
| AMC16 | 63.5 | M | 25 | 12.3 | 14.0 |
| AMC17 | 53 | M | 28 | 14.6 | 13.2 |
| AMC18 | 70 | M | 25 | 9.6 | 10.4 |
| AMC19 | 66 | M | 21 | 11.4 | 9.2 |
| AMC20 | 53 | F | 29 | 17.6 | 10.8 |
| AMC21 | 49 | F | 25 | 14.0 | 11.6 |
| AMC22 | 64 | M | 24 | 13.6 | 10.7 |
| AMC23 | 70 | M | 23 | 12.8 | 12.0 |
| **Mean (SD)** | **62.7 (6.9)** | **15M:8F** | **24.5 (2.3)** | **12.6 (2.3)** | **12.7 (3.4)** |

Abbreviations: MoCA: Montreal cognitive assessment (range 0 to 30, higher scores reflect better cognition, scores $\geq 26$ are considered normal cognition); TUG1: timed up and go test, visit 1; TUG2: timed up and go test, visit 2; AMC: age-matched control.

**Table 3:**
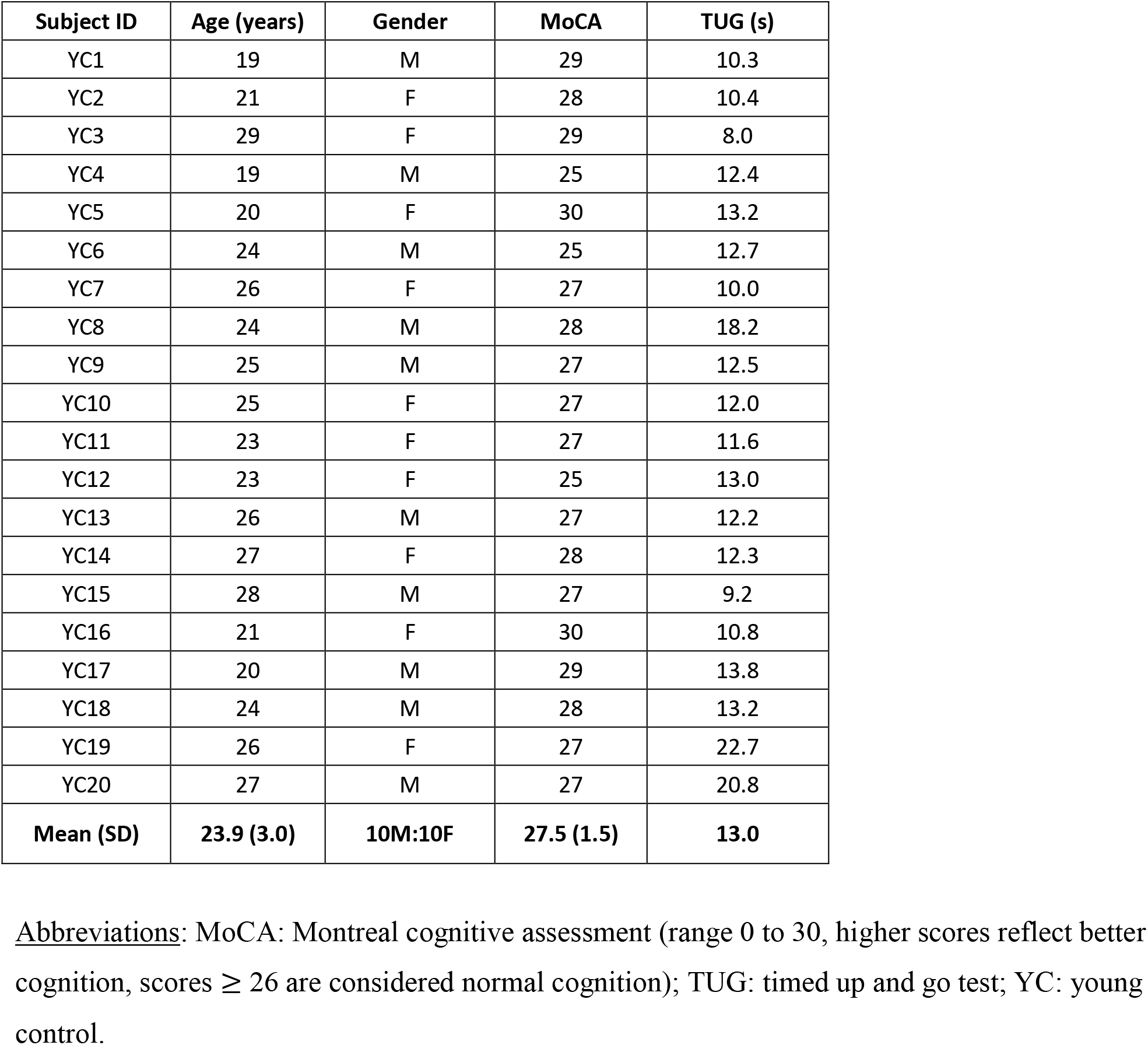
Young control participants’ details.

PD participants and age-matched controls performed the same experiment twice (on two separate visits, one week apart). The primary aim of this was to test for a PD medication effect. Although it would have been preferable to randomly counterbalance on and off medication states across the two visits, many PD patients requested to first experience and perform the experiment in the “on-state”. Hence, for the sake of patient comfort and consistency, we tested all PD patients first in the “on-state”, and a week later, in the “off-state” (at least 12 hours since their last anti-Parkinson medication dose). Accordingly, for better comparison, the age-matched control participants also performed the same experiment twice. Young controls performed the experiment only once. All PD and all but one age-matched participant returned for the second visit.

Gait function was assessed with a “timed up and go” (TUG) test using a smartphone application (Madhushri *et al.*, 2016). TUG results are presented in the respective group tables (Tables 1-3). All participants performed the TUG test twice in succession, prior to the psychophysics task on each visit (TUG scores reflect the average, per visit). PD subjects performed the TUG test more slowly than age-matched controls (14.9 ± 4.9s versus 12.6 ± 2.4s on the first visit and 14.0 ± 2.7s versus 12.7 ± 3.4s on the second visit, F_(1,75)_ 5.6, *p* = 0.02), with no significant effect of visit or visit-group interaction (consistent with the early stage of PD in our cohort). We therefore did not find TUG results useful for further analyses (we used PD UPDRS scores to correlate task performance with disease severity).

### Stimuli and task

The experiments were run in the motion simulator at the Gonda Brain Research center, at Bar Ilan University. Vestibular (inertial motion) stimuli were generated using a six-degrees-of-freedom motion platform (MB-E-6DOF/12/1000KG; Moog Inc.), upon which a car seat was mounted (Fig. 1A). Participants were seated comfortably in the car seat and restrained safely with a four-point seatbelt. Although additional somatosensory or proprioceptive cues may also be used during inertial motion (e.g., cutaneous sensation and muscle proprioception), we refer to this condition as “vestibular” because performance strongly depends on intact vestibular labyrinths (Gu *et al.*, 2007). In addition, even if considered collectively a vestibular-somatosensory-proprioceptive (non-visual) cue, the same rules of cue integration still apply when testing integration with visual cues.

**Figure 1.**
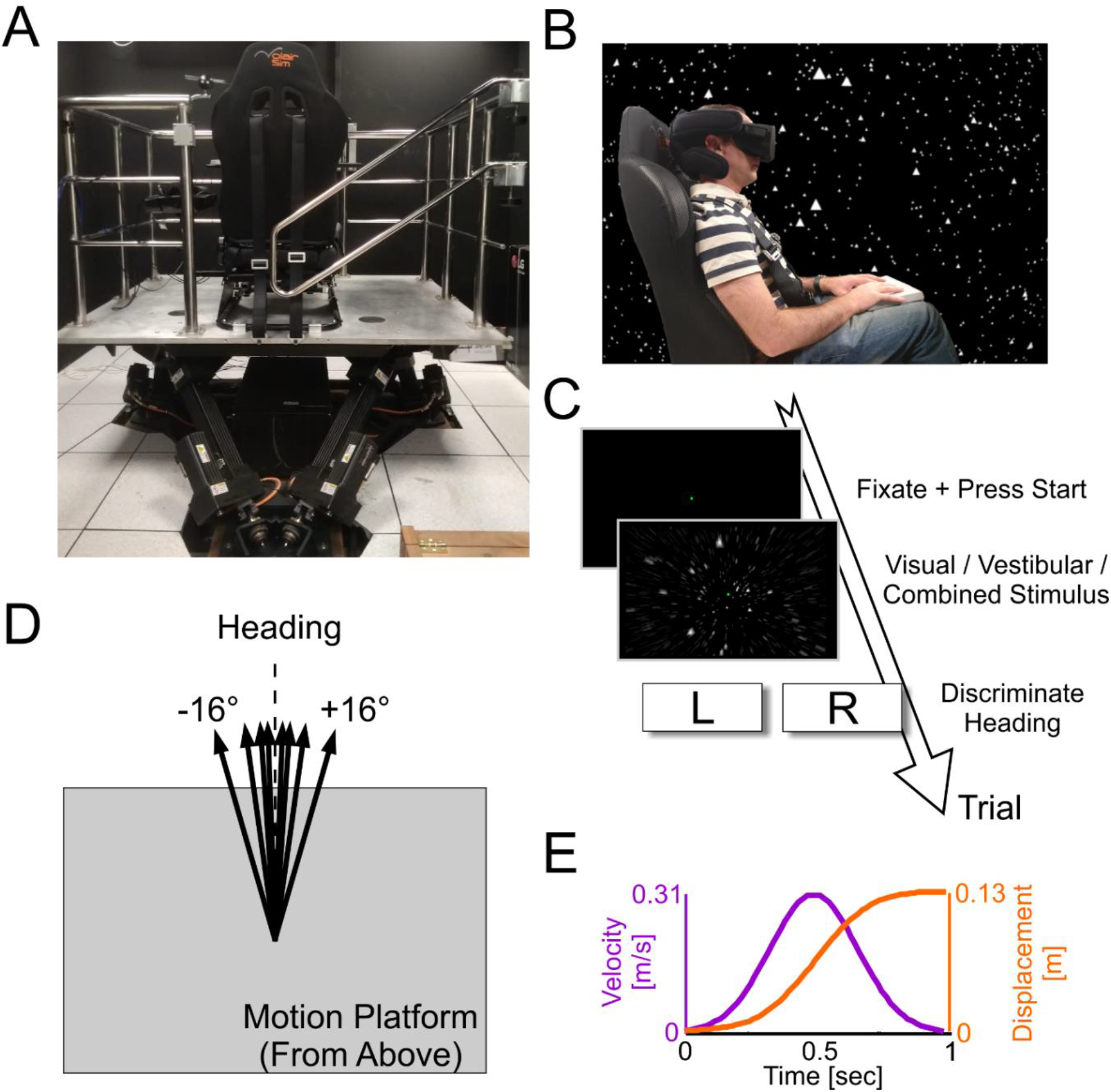
Experimental setup. A) The 6-degrees-of-freedom motion platform, with mounted chair (viewed from behind). B) A participant sitting in the chair (viewed from the side) wearing a head-mounted display (Oculus Rift) for visual optic flow stimuli. The background stars depict what the participant sees in virtual reality (shown here for illustrative purposes only). C) Flow of a single trial: after trial initiation, a 1s (visual, vestibular or combined visual-vestibular) motion stimulus is experienced, after which the participant reports his/her heading discrimination (left or right). D) Schematic of various heading directions. E) Motion profile (the same for all stimuli; only heading direction was varied).

The visual stimulus simulated self-motion through a 3D cloud of dots, and was presented via a virtual reality head-mounted display (HMD, Oculus Rift CV1; Fig. 1B). The star field in the background of Figure 1B reflects the visual experience of the participant in virtual reality. Participants with corrected vision either wore their own glasses (or contact lenses) under the HMD, or we inserted prescription lenses to match the participant’s prescription, from a set made specifically for the Oculus Rift (VR Lens Lab). The participant’s head was supported by a head support with lateral extensions to limit head movement (Black Bear; Matrix Seating Ltd.). Participants wore the HMD, which covered their field of vision, throughout the experiment, and the room was kept dark to avoid any other visual cues.

The (vestibular and visual) self-motion stimuli followed a linear path trajectory (0.13m displacement) in the horizontal plane. These were primarily in the forward direction, but with slight deviations to the right or left of straight ahead (Fig. 1D). Stimulus velocity followed a Gaussian profile (peak velocity 0.31 m/s, and peak acceleration 1.14 m/s^2^), and lasted 1s (Fig. 1E). A single interval stimulus was presented on each trial, which was either unisensory vestibular (inertial motion in darkness), unisensory visual (optic-flow), or multisensory visual-vestibular (inertial motion with simultaneous optic flow). A central fixation point was displayed via the HMD on all trials, rendered at a fixed distance of 0.62m straight in front of the participant. It remained at this location (in relation to the participant) also during the motion stimuli. Participants were instructed to maintain fixation on this point throughout each trial.

Participants held a control box (Cedrus RB-540) which rested on their lap. They initiated trials at their own pace by pressing the center (start) button and reported their perceived heading direction, after the stimulus ended, by pressing the right or left button on the response box. Three different auditory tones were used to indicate: (i) that the system was ready for a new trial (i.e. to press start), (ii) that a choice was registered, and (iii) a response timeout (2s after the end of the stimulus, if a choice was not registered). Participants were instructed to avoid this time-out by making a timely response on every trial, and to guess when unsure. All participants underwent brief training with practice trials and verbal feedback from the experimenter in order to confirm that they understood the instructions and performed the task adequately before starting the actual experiment. An intercom system enabled the participants and the experimenter to communicate throughout the duration of the experiment. No feedback was provided during the experiment regarding whether their answer was correct/incorrect.

Possible stimulus heading values were distributed logarithmically around straight ahead at angles of ±16°, 8°, 4°, 2°, 1°, 0.5° or 0.25°, where zero represents straight ahead and positive (or negative) values represent a rightward (or leftward) deviation from straight ahead. The absolute heading magnitude was set according to a staircase procedure (Cornsweet, 1962) and the heading sign (positive or negative) was selected randomly for each trial. A separate staircase procedure was run per condition (with pseudo-randomly interleaved trials). Each staircase began with the easiest heading (±16°). After a correct response, heading magnitude was reduced (such that the task became more difficult) 30% of the time, and remained unchanged 70% of the time. After an incorrect response, it was increased (such that the task became easier) 80% of the time, and remained unchanged 20% of the time. This staircase rule converges at ~73% correct responses, thereby sampling an information-rich region of the psychometric function, on an individual basis.

Visual cue reliability was controlled by manipulating visual motion coherence. Two coherence levels were used to test visual performance: (1) 100% coherence, in which all the dots moved coherently according to the direction of simulated self-motion, and (2) 65% coherence, in which 65% of the dots moved coherently, and the remaining 35% moved randomly. We chose 65% coherence because multisensory integration is best studied when the unisensory (visual and vestibular) cue reliabilities are similar (Angelaki et al., 2009) and pilot experiments in our system showed that visual reliability at 65% coherence was roughly similar to vestibular performance. Accordingly, multisensory conditions were also tested at 65% coherence. Unisensory visual performance was also tested at 100% coherence, in order to disambiguate whether any observed visual deficit resulted from impaired visual perception per se, or from sensitivity to sensory noise Zaidel *et al.*, (2015). Vestibular reliability was not manipulated.

To test and quantify multisensory cue weighting, a slight discrepancy (Δ) between the visual and vestibular headings was introduced when they were presented in combination. By convention, Δ > 0 means that the vestibular headings were offset to the right, and the visual headings were offset to the left, each by Δ/2 (and vice-versa for Δ < 0). Having a set discrepancy between the visual and vestibular headings allowed us to measure the relative cue weighting. For example, for evenly weighted cues, the multisensory percept should lie exactly in between the two. By contrast, when one cue is dominant, then the multisensory percept should lie closer to the dominant cue (this is explained further, quantitatively, below).

Five stimulus conditions were run (interleaved): (i) vestibular-only, (ii) visual-only with 100% visual coherence, (iii) visual-only with 65% visual coherence, (iv) combined visual-vestibular with 65% visual coherence and Δ = +6°, and (v) combined visual-vestibular with 65% visual coherence and Δ = −6°. Δ = ±6° was used, since it is well within the range of values that are integrated despite the discrepancy (Acerbi *et al.*, 2018). A total of 400 trials were collected per visit (80 trials for each of the 5 stimulus conditions). The experiment was divided into two blocks of 200 trials, approximately 20 minutes per block, to allow the participants to take a break in the middle.

### Data analyses and statistics

Data analyses were performed with custom software using Matlab R2013b (MathWorks) and *psignifit* toolbox for Matlab version 4 (Schütt *et al.*, 2016). Psychometric plots represent the proportion of rightward choices as a function of heading angle and were calculated by fitting the data with a cumulative Gaussian distribution function (see examples in Fig. 2). Separate psychometric functions were fit per participant for each stimulus condition. The mean (µ) of the fitted cumulative Gaussian distribution function represents the point of subjective equality (PSE; namely, the heading for which the probability of choosing right or left is *p* = 0.5). And, the psychophysical ‘threshold’ is defined as the standard deviation (σ) of the fitted cumulative Gaussian distribution function. Lower threshold values reflect better (more precise) performance. Since thresholds are nonnegative values that scale geometrically, logarithmic values were used for statistics and plotting. For better threshold estimation, lapse rates were also simultaneously fit (with a narrow prior, up to 0.1) and the thresholds’ prior was extended to allow for high threshold values amongst some of the participants.

**Figure 2.**
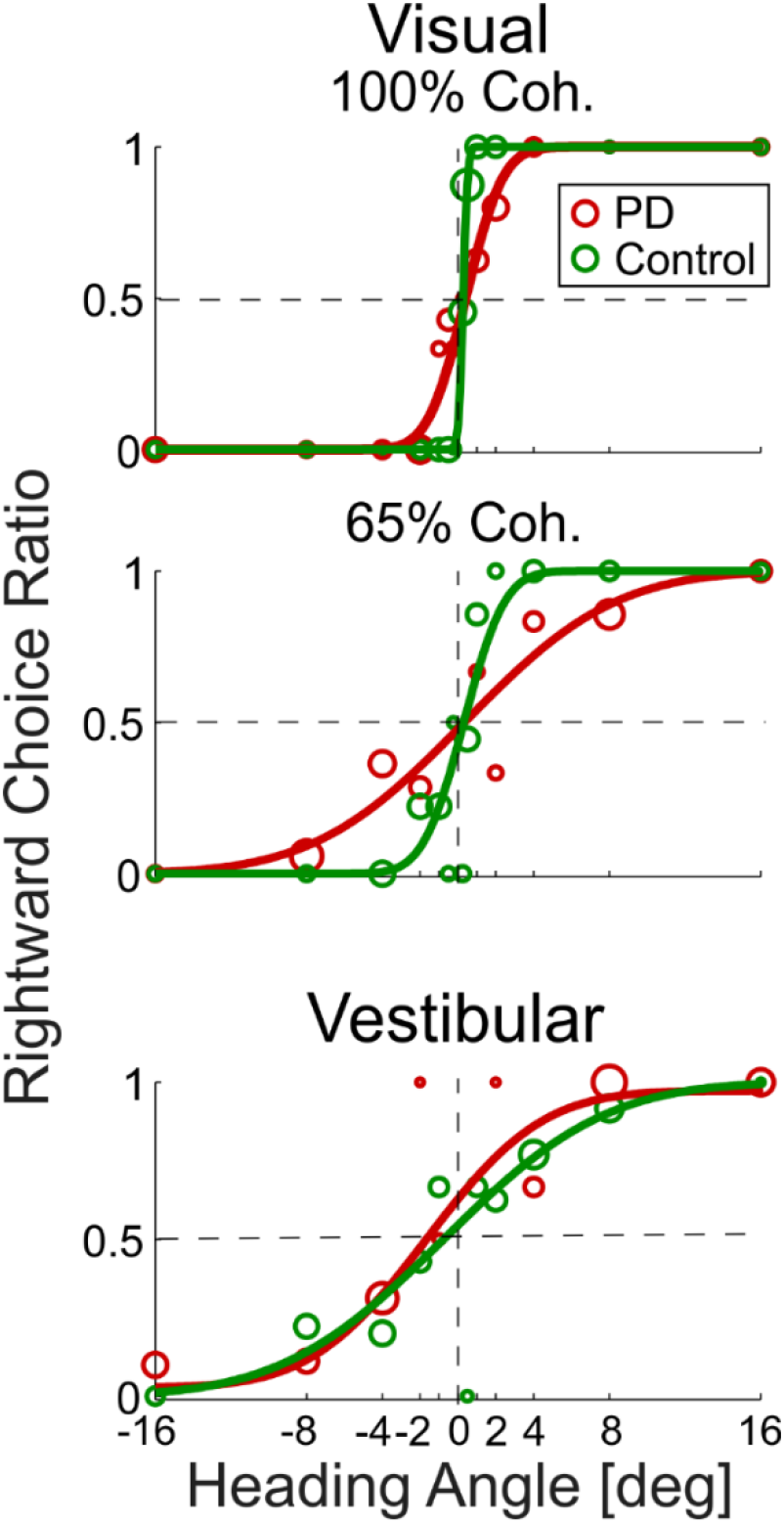
Example psychometric plots. Behavioral responses to visual stimuli at 100% coherence (top plot), 65% coherence (middle plot) and vestibular stimuli (bottom plot) are presented for an example Parkinson’s disease (PD, red) and an example age-matched control participant (green). Circle markers represent the ratio of rightward choices for a specific heading (marker size reflects the number of trials collected at that heading). The data were fitted with cumulative Gaussian distribution functions (solid lines). The vertical dashed line marks heading = 0°, and the horizontal dashed line marks y = 0.5 (equal probability of rightward and leftward choices).

Statistical analyses were performed using JASP (version 0.11.1) and Matlab. To compare visual thresholds, we applied a three-way repeated measures ANOVA defining visual coherence (two levels: 100% and 65%) and visit (Visit 1 and Visit 2) as within-subject factors, and group (PD and age-matched controls) as the between-subject factor. The same was done for vestibular thresholds, but without coherence as a factor. This allowed us to test for both main effects and interactions. The young controls were not part of those ANOVA comparisons because they were only tested in one visit (and the primary comparison in this study was PD vs. age-matched controls). Hence, further analyses, per visit, were also performed. Additional details of these specific statistical comparisons are presented together with the results below.

### Bayesian multisensory integration

The Bayesian framework for multisensory integration has been well-described previously. See Angelaki et al. (2009) for review. We briefly summarize it here, as it relates to our study. When presented with a visual or vestibular heading stimulus (*s*_*vis*_ or *s*_*ves*_, respectively) the participant will have a noisy internal measurement of that stimulus (*x*_*vis*_ or *x*_*ves*_, respectively). These measurements are assumed to be normally distributed around the stimulus: 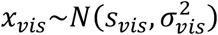 and 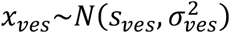, where 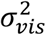 and 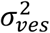 represent visual and vestibular noise distribution variances, respectively. Cue reliability (*R*) is defined as the inverse variance:

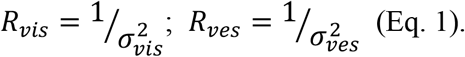

Thus visual and vestibular cue reliabilities can be estimated from the unisensory thresholds (taken from the standard deviations, *σ*_*vis*_ or *σ*_*ves*_, of the fitted Gaussian psychometric curves; see section **Data analyses and statistics**).

In the multisensory condition, when both visual and vestibular measurements are attained, the optimal estimate of the stimulus (assuming a flat prior) is a linear weighted combination of the measurements:

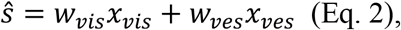

where the cue weights:

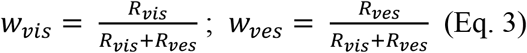

reflect their relative reliabilities (note that the weights sum to 1; *w*_*vis*_ + *w*_*ves*_ = 1). Intuitively, when *σ*_*vis*_ < *σ*_*ves*_ the visual cue is relatively more reliable (*R*_*vis*_ > *R*_*ves*_). Hence, the visual measurement should be given more weight than the vestibular measurement, during multisensory integration.

To test whether PD patients indeed followed Bayesian optimal cue weighting, we compared the Bayesian predicted weights (Eq. 3) to the actual (empirical, observed) visual and vestibular weights, estimated from the multisensory conditions. In order to estimate these actual weights, a systematic discrepancy (Δ = *s*_*ves*_ − *s*_*vis*_) was introduced between the cues in the multisensory condition (Δ = ±6° in our study). The actual visual weights could then be estimated as follows (Fetsch *et al.*, 2009):

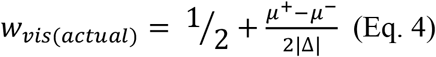

Where µ^+^ and µ^−^ are the PSEs of the combined cue conditions with positive and negative Δ, respectively. The vestibular weights are then calculated by:

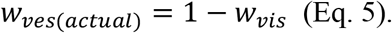

Bayesian optimal integration also predicts that the multisensory (combined) cue threshold, should be lower than, and can be quantitatively predicted from, the unisensory thresholds (Ma *et al.*, 2006) as follows:

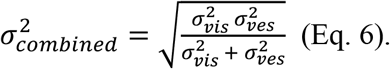

We also compared the Bayesian prediction for the combined thresholds to the actual (empirical) combined cue thresholds measured from the multisensory conditions. Since there were two combined conditions (Δ = ±6) the (geometric) mean of the two thresholds was used.

### Data availability

The data are available upon request from the authors.

## RESULTS

In this study, we tested visual and vestibular (unisensory) self-motion perception, as well as visual-vestibular multisensory integration, in PD patients. We found impaired visual self-motion perception in PD, but normal vestibular self-motion perception. Furthermore, in the multisensory condition, PD patients over-weighted their (impaired) visual cues, deviating from Bayesian predictions of optimal integration. Details and expansion of these results are presented below.

### Impaired visual self-motion perception in PD

Unisensory visual and vestibular performance of an example PD patient and an example age-matched control are presented in Figure 2. A steeper psychometric curve (one that approaches a step function) reflects better performance – namely, higher precision in discriminating rightward from leftward heading stimuli. While poorer performance is marked by a flatter (less steep) curve. In Figure 2, the example PD participant (red) demonstrates worse visual self-motion perception (flatter psychometric curve) compared to the control participant (green) – for both 100% and 65% visual motion coherence conditions (top and middle plots, respectively). By contrast, they had similar vestibular self-motion perception (bottom plot). To compare performance quantitatively, and across groups, perceptual thresholds were measured from the psychometric data, per participant, condition and visit (see Methods).

PD patients had consistently higher visual thresholds (i.e., worse visual performance) in all conditions (Fig. 3, left column) – the red line (depicting visual thresholds in PD) lies above the others for both 100% and 65% coherence levels, and for both the first (Fig. 3A) and the second (Fig. 3B) visit. Comparing PD to age-matched controls revealed that the increase in PD visual thresholds is significant (*F*_(1,39)_ = 4.9, *p* = 0.03, repeated measures three-way ANOVA: 2 groups × 2 coherences × 2 visits; young controls performed the task on one visit only, and are therefore compared in a separate analysis below). A significant effect of coherence (*F*_(1,39)_ = 60.3, *p* < 0.001) trivially reflects the manipulation of visual coherence (higher thresholds for 65% vs. 100% coherence). No significant interactions (between group, coherence and visit) were found (*p* > 0.16). These results therefore indicate that visual self-motion perception is generally impaired in PD.

**Figure 3.**
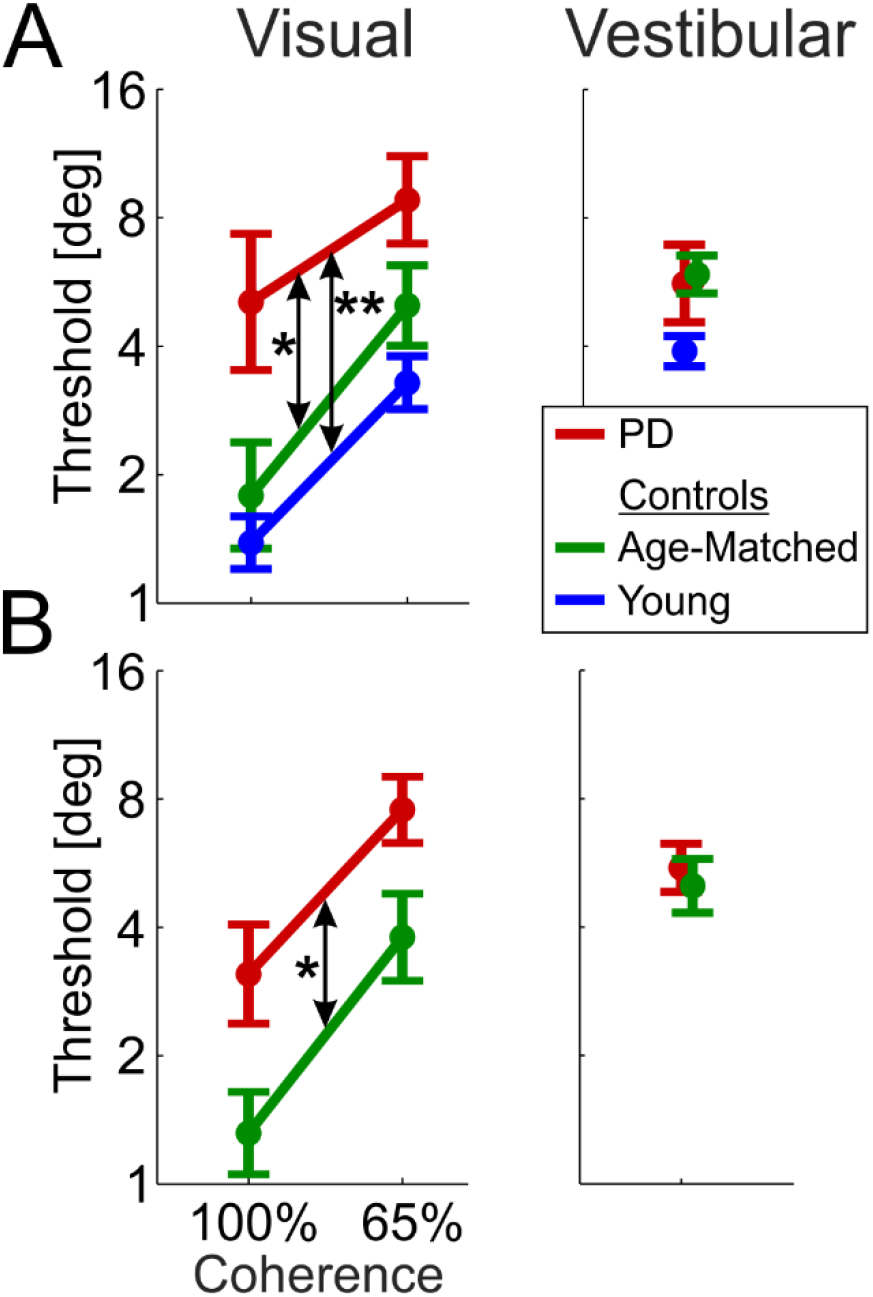
Impaired visual self-motion perception in PD. Visual and vestibular thresholds (left and right columns, respectively) are presented for the Parkinson’s disease (PD, red), age-matched (green) and young control (blue) groups. A) Data from Visit 1 (PD on medication). B) Data from Visit 2 (PD off medication). Young controls were tested only once (Visit 1). Data points and error bars represent mean ± SEM. * indicates *p* < 0.05; ** indicates *p* < 0.01.

Further analysis, per visit, showed that PD patients had significantly impaired visual performance on both visits, independently. On the first visit PD visual thresholds were significantly higher vs. both age-matched and young control groups (*p* = 0.04 and *p* = 0.003, respectively, post-hoc comparison with Tukey correction; repeated measures two-way ANOVA: 3 groups × 2 coherences; Fig. 3A). Young healthy controls had slightly lower visual thresholds than age-matched controls (but this was not significant, *p*_tukey_ = 0.55). Also on the second visit, PD visual thresholds were significantly higher vs. age-matched controls (*F*_(1,39)_ = 4.3, *p* = 0.045; repeated measures two-way ANOVA: 2 groups × 2 coherences; Fig. 3B). Finding the same result, of increased visual thresholds in PD, on both visits (independently), strengthens the finding.

By contrast, vestibular thresholds for the PD and age-matched control groups were highly overlapping in both the first and the second visit (red and green, respectively; Fig. 3, right column) and did not differ statistically (*F*_(1,39)_ = 0.033, *p* = 0.86; repeated measures two-way ANOVA: 2 groups × 2 visits). This indicates that vestibular self-motion perception was not impaired in the PD group. Although there was a trend for lower vestibular thresholds in the young controls (first visit; Fig 3A, right), this did not reach significance (*F*_(2,59)_ = 2.74, *p* = 0.07; one-way ANOVA: 3 groups). Comparable performance of PD and age-matched controls in the vestibular condition indicates that impaired performance in the visual condition did not arise from other difficulties in task performance (e.g., reporting choices) and validates the use of a 2AFC task to probe perceptual function in PD. The stark difference between the visual and vestibular results (significantly impaired visual, but intact vestibular performance in PD) tested in the same (interleaved) task, with the same participants, points to a specific impairment of visual self-motion perception in PD.

One might have expected worse performance in the second vs. the first visit in the PD group (off vs. on medication). But this was not observed. Rather, both groups had improved visual thresholds on the second visit (*p* = 0.02, three-way ANOVA presented above, Fig. 3 left) without any group × visit interaction (*p* = 0.93). Vestibular thresholds were unchanged for both groups (*p* = 0.4, two-way ANOVA presented above, Fig. 3 right) also with no group × visit interaction (*p* = 0.53). These results suggest that PD patients responded to the second visit in the same way as age-matched controls (with a small learning effect for visual cues) and that medication status on/off did not improve or impair self-motion perception. Additional studies, with counterbalanced medication state across visits (and larger sample size) might uncover subtle and specific medication effects.

### Visual thresholds correlate with motor impairment

To further investigate the relationship between impaired visual self-motion perception and PD, we tested whether perceptual thresholds correlate with disease severity (motor UPDRS scores). For visual thresholds, four (Pearson) correlations were tested (2 coherence levels × 2 visits). UPDRS scores on and off medication were used for the first and the second visit, respectively. All four correlations were statistically significant: *r* = 0.51, *p* = 0.026 and *r* = 0.63, *p* = 0.004 (100% and 65% coherence, respectively, for the first visit; Fig. 4A, left), and *r* = 0.69, *p* = 0.002 and *r* = 0.60, *p* = 0.007 (100% and 65% coherence, respectively, for the second visit; Fig. 4B, left). By contrast, no significant correlation was seen between vestibular thresholds and UPDRS scores (*r* = 0.34, *p* = 0.15 and *r* = 0.05, *p* = 0.84 for the first and the second visit, respectively; Fig 4, right column).

**Figure 4.**
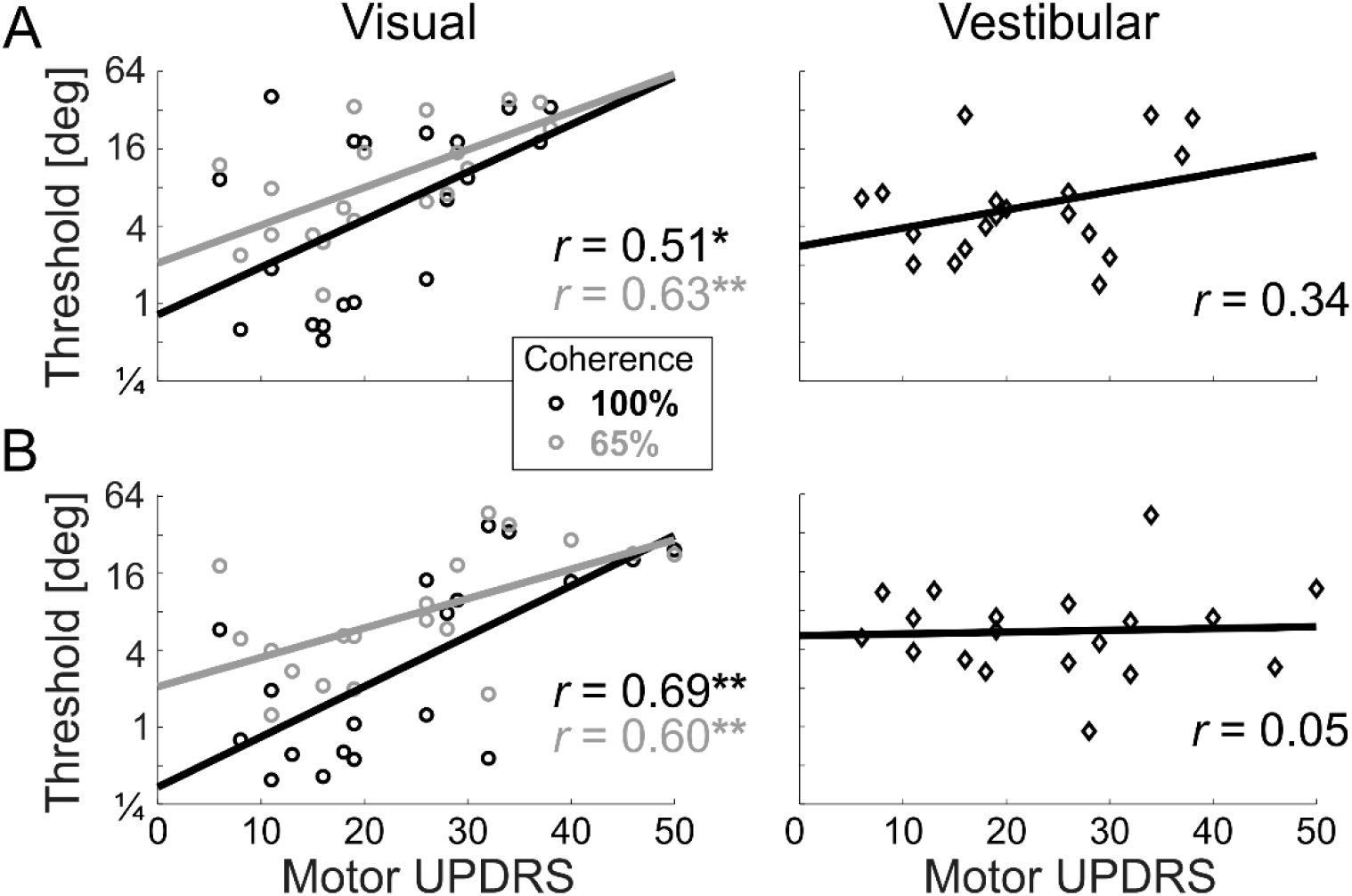
Visual self-motion perception deteriorates with PD severity. For the Parkinson’s disease (PD) participants, visual and vestibular thresholds (left and right columns, respectively) are presented vs. their Unified Parkinson’s disease rating scale (UPDRS) motor scores. A) Data from Visit 1 (on medication). B) Data from Visit 2 (off medication). Black and gray circle markers represent visual thresholds at 100% and 65% coherence, respectively. Black diamonds present vestibular thresholds. Solid lines and ‘r’ values represent the linear regressions and correlation values of the respective plots. * indicates p < 0.05; ** indicates p < 0.01.

Thus, visual (but not vestibular) self-motion perception deteriorates with disease severity. Furthermore, the correlation scores (*r* = 0.51 – 0.69) indicate that between 26% - 48% of the variance in visual thresholds can be attributed to PD disease severity. This is striking in light of the high variance of perceptual thresholds typically observed across individuals. Although task performance could also correlate with cognitive function, this would not explain any of our results since the PD and age-matched groups had comparable scores (presented above and in Tables 1-2). We further confirmed this by adding MoCA as a covariate to the three-way ANOVA presented above. We found that also when controlling for cognitive function, PD visual self-motion performance was significantly impaired vs. age matched controls (*p* = 0.012). A trend for lower visual thresholds with better MoCA scores was seen, but this was not significant (*p* = 0.06).

### Visual overweighting in PD

Our second main aim in this study (beyond testing unisensory visual and vestibular self-motion perception in PD) was to examine how PD patients integrate information from visual and vestibular cues. Bayesian theory of multisensory integration provides two specific testable predictions of optimal integration: (i) the weights attributed by a participant to each cue, during integration, should equal the relative reliabilities of the respective cues, such that the more reliable cue is more heavily weighted. Quantitatively, the predicted weights can be calculated from the unisensory thresholds (using Equations 1 and 3). And these can be compared to the actual (empirically observed) cue weights, measured from the multisensory conditions (using Equations 4 and 5). (ii) A participant’s multisensory (combined cue) threshold should be lower (better) than each of his/her unisensory thresholds. Also multisensory thresholds can be quantitatively predicted from the unisensory thresholds (using Equation 6) and compared to the actual (empirically observed) multisensory thresholds. Thresholds (and thus predicted weights) are personal quantities, which depend on each individual’s function. Therefore, these two predictions were tested per participant.

For the first prediction (regarding cue weighting), it is sufficient to compare empirically observed vs. predicted weights for one cue (cue weights sum to one, so results for the second cue are complementary). Here, we present this comparison for visual weights (from the first and the second visit in Fig. 5A and B, respectively). Scatter plots (Fig. 5, left column) depict observed vs. predicted visual weights for PD, age-matched, and young participants (red, green and blue ‘o’ markers, respectively) as well as group means ± SEM (respectively colored ‘+’ markers). Deviations from the diagonal black lines (which represent perfect predictions) to the upper left (or lower right) indicate visual (or vestibular) overweighting. These deviations were analyzed, per group and visit, using two-tailed paired *t*-tests (with Bonferroni correction for five comparisons) and compared between PD and age-matched controls across visits using a repeated measures ANOVA.

**Figure 5.**
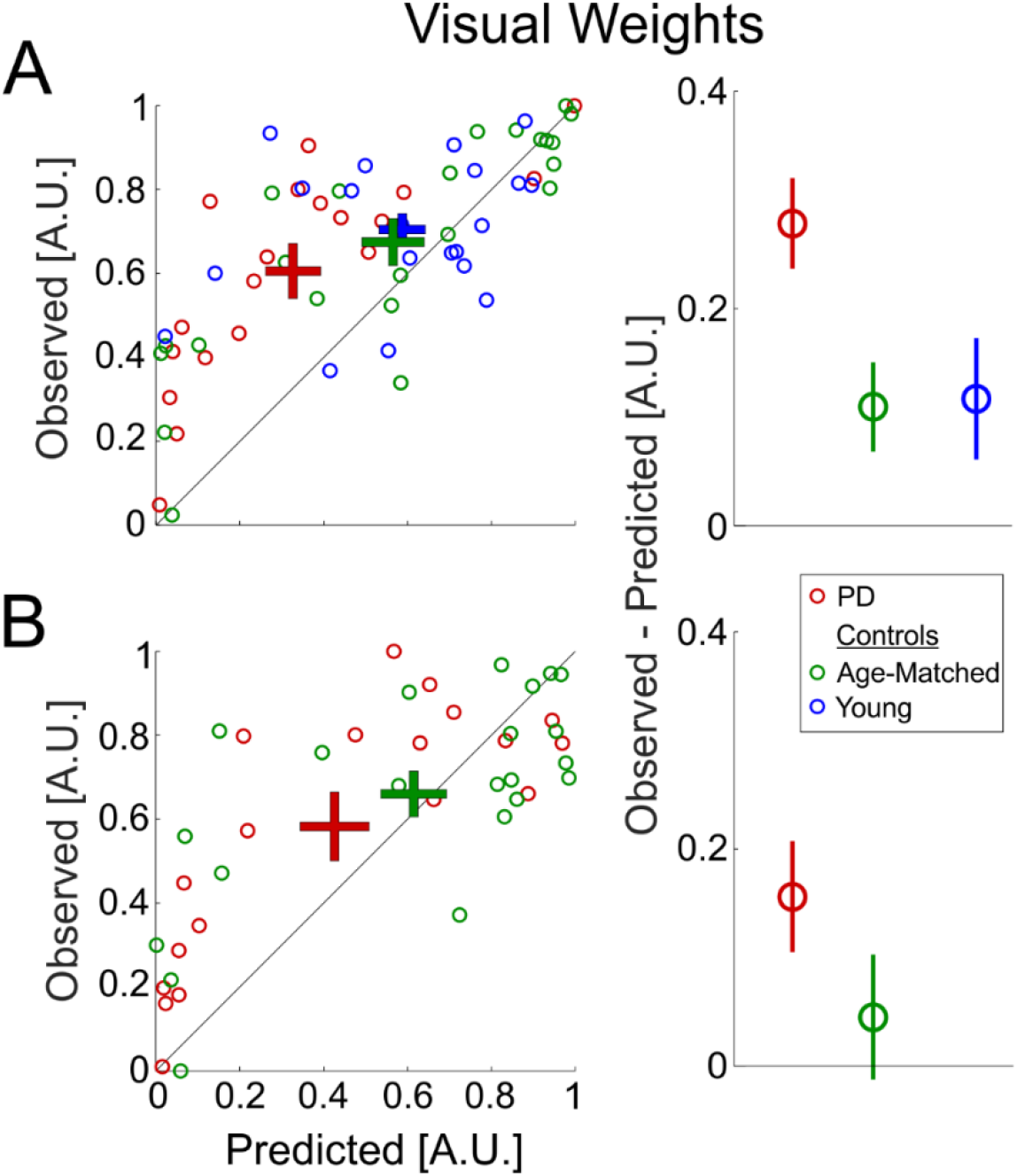
Visual overweighting in PD. A) Data from Visit 1 (PD on medication). B) Data from Visit 2 (PD off medication). Young controls were tested only once (Visit 1). Left column (scatter plot): each data point depicts the observed visual weight of an individual participant (extracted from the combined cue Δ conditions) vs. the Bayesian predicted visual weights (estimated from the unisensory conditions). ‘+’ markers represent the mean ± SEM for each group (by respective color). The diagonal black line (y = x) represents equality between observed and predicted weights. Right column: the difference between the observed and predicted visual weights (mean ± SEM) for each group (by respective color).

PD patients had the largest deviations from optimality, with significant visual over-weighting, on both the first and the second visit (*t*_(18)_ = 6.7, *p* = 10^−5^ and *t*_(18)_ = 3.14, *p* = 0.028, respectively). Although visual overweighting was also seen for the control groups, this was to a lesser degree, and did not reach significance (*t*_(22)_ = 2.66, *p* = 0.07 and *t*_(19)_ = 2.09, *p* = 0.25 for age-matched and young controls, respectively, on the first visit, and *t*_(21)_ = 0.78, *p >* 1 for age-matched controls on the second visit). The difference between the observed and predicted weights (Fig. 5, right column) was significantly larger in PD vs. age-matched controls (*F*_(1,39)_ = 5.1, *p* = 0.03; repeated measures two-way ANOVA: 2 groups × 2 visits) indicating greater visual over-weighting in PD. Visual overweighting was reduced for both groups in the second visit (*F*_(1,39)_ = 8.4, *p* = 0.006) with no significant group × visit interaction (*F*_(1,39)_ = 0.6, *p* = 0.4). This indicates a possible practice effect for both PD and age-matched controls (in line with improved visual thresholds, described above). And, like above, no evidence for a medication effect.

The second Bayesian prediction is a little more difficult to discern, since the largest expected reduction of multisensory (vs. unisensory) thresholds is only by a factor of 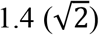 and occurs when the unisensory thresholds are equal (Eq. 6). Visual (65% coherence) and vestibular thresholds were indeed roughly similar per group, but less so for the PD group, who had larger visual vs. vestibular thresholds (Fig. 6, right column) making it more difficult to discern an integration deficit. A trend was seen for larger observed vs. predicted thresholds in PD on their first visit (Fig. 6A, right plot, and red ‘+’ marker lying above the diagonal line in Fig. 6A, left plot), however, this was not significant (*p* = 0.33 after Bonferroni correction for 5 comparisons, paired *t*-test). Multisensory thresholds for the two control groups on the first visit were consistent with optimal integration (blue and green ‘+’ markers lying close to the diagonal in Fig. 6A, left plot, and comparable observed and predicted multisensory thresholds in Fig. 6A, right plot; *p* > 1 after Bonferroni correction, paired *t*-tests). Also on the second visit there were no significant differences between observed and predicted thresholds, for both groups (p > 0.16 after Bonferroni correction, paired *t*-tests).

**Figure 6.**
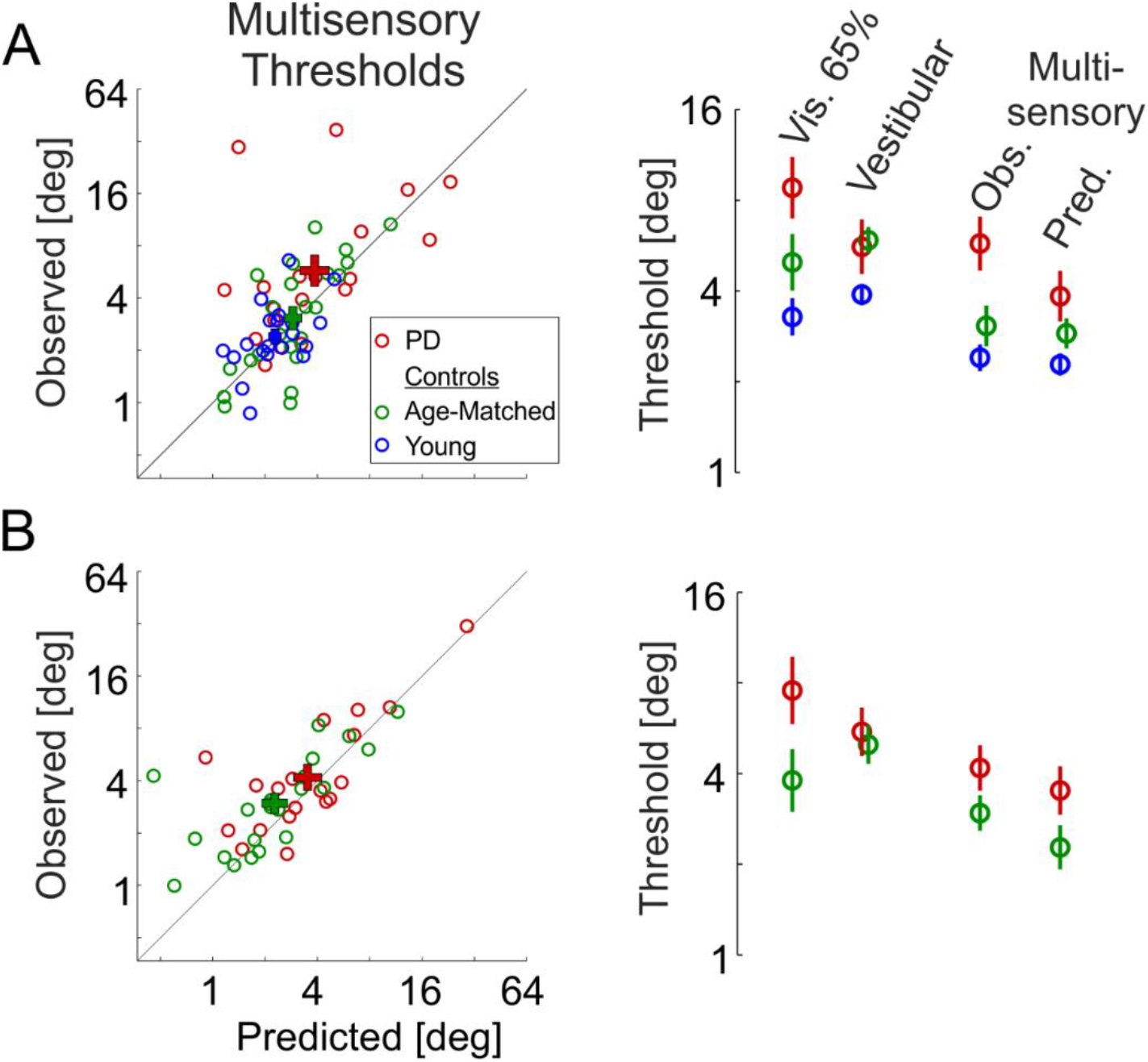
Multisensory thresholds. A) Data from Visit 1 (PD on medication). B) Data from Visit 2 (PD off medication). Young controls were tested only once (Visit 1). Left column: each data point depicts the observed combined cue (multisensory) threshold of an individual participant vs. the Bayesian prediction (estimated from the unisensory conditions). ‘+’ markers represent the mean ± SEM for each group (by respective color). The diagonal black line (y = x) represents equality between observed and predicted thresholds. Right column: mean ± SEM unisensory (visual 65% coherence and vestibular) and multisensory (observed and predicted) thresholds, per group (by respective color).

## DISCUSSION

In this study we investigated perception of self-motion in PD. To measure unisensory visual and vestibular function, separately, and to test multisensory integration of visual and vestibular cues, we used a well-established psychophysics paradigm that has not been previously applied to study PD. Advantages of this paradigm are: i) it is not dependent on graded motor responses (which are themselves altered in PD). Rather, it relies on categorical choices (right or left), which are analyzed based on signal detection theory to provide a canonical metric for perceptual function. ii) It allowed us to quantify multisensory integration within the principled framework of Bayesian inference.

The study results provide several novel and interesting findings: (1) PD patients demonstrated impaired visual self-motion perception, which deteriorates with PD severity. (2) PD vestibular function was indistinguishable from age-matched controls. (3) PD patients over-weighted visual (vs. vestibular) cues during multisensory integration. Together these results indicate that self-motion perception is affected in PD both by impaired visual cues, and by suboptimal multisensory integration. We further discuss these results and their implications here below.

Altered veering in response to visual optic flow has been described in PD (Davidsdottir *et al.*, 2008; Lin *et al.*, 2014). However, with graded motor actions (veering) as responses, those studies do not dissociate perceptual from motor or sensorimotor dysfunction. Here we found a specific impairment of visual self-motion perception in PD. This does not reflect a general deficit of visual motion perception, since the ability to discriminate aggregate motion of a (flat) random dot kinematogram (RDK; dots moving on a 2D screen; not simulating self-motion) is not impaired in PD (Putcha *et al.*, 2014; Jaywant *et al.*, 2016). Thus, visual self-motion perception (a skill which is vital for proficient balance and gait) seems specifically impaired in PD. This might reflect the higher complexity of perceptual processing required to differentiate one’s own motion from motion of objects in the environment (Dokka *et al.*, 2015) and/or the 3D nature of our stimuli and experiment.

Our concurrent observation of unimpaired performance with vestibular stimuli (in the same participants and task) firstly confirms that impaired visual performance was not due to difficulty in performing the task itself, but rather reflects a specific impairment of visual self-motion perception. Notably, it also suggests that vestibular performance seems largely spared in PD. This is in line with Bertolini et al. (2015) who also did not find vestibular sensory impairment (but rather found a central vestibular integration failure). Accordingly, altered nystagmus in PD to caloric stimulation (Reichert *et al.*, 1982) might reflect sensory-motor integration deficits, rather than vestibular deficits per se. However, it is also possible that vestibular impairment might only emerge at a later stage in PD (our cohort was early stage). Nonetheless, the stark difference between the visual and vestibular results suggests a stronger visual self-motion impairment, seen already at an early stage.

Humans (and animals) have been shown, in many tasks and across different modalities, to integrate their senses in a near-optimal Bayesian manner (Jacobs, 1999; Landy and Kojima, 2001; Ernst and Banks, 2002; Alais and Burr, 2004; Butler *et al.*, 2010; Raposo *et al.*, 2012). Thus, visual over-weighting to the extent that we find here in PD, is quite rare. Furthermore, in this same visual-vestibular self-motion task, humans (and monkeys) have been shown to dynamically reweight visual and vestibular cues in accordance with relative cue reliability, even on single trials (Fetsch *et al.*, 2009). Yet, despite visual self-motion perception presumably deteriorating in PD over a long time, their integration weights seems to reflect their original, unimpaired state.

There are several possible explanations to this finding, which may not be mutually exclusive. (1) It could reflect a general deficit in multisensory integration in PD, perhaps related to general cognitive inflexibility (Cools *et al.*, 2001). (2) It may reflect a specific over-estimation of visual cue reliability in PD (i.e., not correctly estimating the current state of visual function) which would lead to visual overweighting. In support of this idea, PD patients seem to demonstrate increased visual dependence (Cooke *et al.*, 1978; Azulay *et al.*, 2002; Vaugoyeau *et al.*, 2007; Davidsdottir *et al.*, 2008; Barnett-Cowan *et al.*, 2010; Funato *et al.*, 2010) despite their many visual deficits (see Weil *et al.*, 2016 for review). Also, PD patients (with dyskinesia) show increased sensitivity to unreliable visual input (Stevenson *et al.*, 2014). Therefore, it is possible that PD patients specifically overestimate the reliability of their current visual function. Further research is needed to tease apart these options, by testing multisensory integration between other modalities, directly investigating PD estimates of their own visual reliability, and investigating the possible effects of medication on altered multisensory integration.

Perception and action are intricately interconnected (Gibson, 1966; Prinz, 1997; Goodale and Westwood, 2004; Warren, 2006; Merriam *et al.*, 2007; Turvey, 2007). Sensory and perceptual deficits in PD contribute to motor impairment (Konczak *et al.*, 2009), and freezing of gait (Almeida and Lebold, 2010). Conversely, sensory input, such as auditory clicks, or a visual grid of tiles on the floor can aid PD function (Bagley *et al.*, 1991; Freeman *et al.*, 1993; Lim *et al.*, 2005). An important motivation for studying self-motion perception in PD is to gain better insight into the mechanisms of balance and gait dysfunction, which are disabling, difficult to treat, and increase the risk of falling. Here, we did not directly test the connection between impaired self-motion perception and gait and balance disorders. However, a connection is likely, and should be investigated in future studies.

Our results have implications for early detection of PD. Certain non-motor symptoms including sensory, perceptual and cognitive impairments may be present even before the diagnosis of PD. Identification of these impairments in the pre-diagnostic or prodromal phase of the disease would be highly relevant clinically, e.g., for possible neuro-protective treatments (Noyce *et al.*, 2016). In our cohort many (8 out of 19) patients were early in the course of the disease (disease duration <4 years) and most (14 out of 19) were levodopa naïve. It is therefore possible that visual self-motion impairment and/or visual over-weighting could be used as a biomarker that is abnormal at clinical PD diagnosis or possibly even before.

An additional potential clinical application is for PD subtyping. Impaired visual self-motion perception may be associated with certain PD subtypes defined clinically (Lord *et al.*, 2014; Mollenhauer *et al.*, 2014) and/or genetically (Neumann *et al.*, 2009; Alcalay *et al.*, 2012; Wang *et al.*, 2014; Zokaei *et al.*, 2014). We did not find a significant correlation between impaired visual self-motion perception and cognitive function. But, the trend may suggest a possible connection, in line with a relationship between other visual impairments and cognitive decline in PD (Gagnon *et al.*, 2009; Weil *et al.*, 2016).

In summary, we have shown here that PD patients have impaired self-motion perception. This is specifically driven by a deficit in visual self-motion cues. Furthermore, these impaired visual cues of self-motion are over-weighted when integrated with largely intact vestibular cues, leading to suboptimal multisensory integration. Defective self-motion perception and suboptimal multisensory integration may have a profound impact on function in PD, and our understanding thereof.

## ACKNOWLEDGEMENTS

We would like to thank Avraham Elkaras for programming assistance, David Swissa for mechanical assistance and Judith Sonn and Tamar Harpaz for management assistance. We would also like to thank the staff of the Movement Disorders Institute at Sheba Medical Center for help with patient recruitment. This work was supported by grants from the National Institute for Psychobiology in Israel (NIPI, 235-17-18) and The Israeli Centers of Research Excellence (I-CORE, Center No. 51/11) to A.Z.

